# Development of a fast feature extraction method for SARS-CoV-2 spike sequences using amino acid physicochemical properties

**DOI:** 10.1101/2023.11.18.567675

**Authors:** Hiroya Oka, Kosaku Noba, Jun Sasahara, Takayuki Hashimoto, Shogo Yoshimoto, Hirohiko Niioka, Jun Miyake, Katsutoshi Hori

**Affiliations:** Department of Biomolecular Engineering, Graduate School of Engineering, Nagoya University, Furo-cho, Chikusa-ku, Nagoya, Aichi 464-8603, Japan; Graduate School of Information Science and Technology, Osaka University, 1-5, Yamadaoka, Suita, Osaka 565-0871, Japan; Graduate School of Engineering, Osaka University, 2-1, Yamadaoka, Suita, Osaka 586-0871, Japan

## Abstract

COVID-19 continues to spread today, leading to an accumulation of SARS-CoV-2 virus mutations in databases, and large amounts of genomic datasets are currently available. However, due to these large datasets, utilizing this amount of sequence data without random sampling is challenging. Major difficulties for downstream analyses include the increase in the dimension size along with the conversion of sequences into numerical values when using conventional amino acid representation methods, such as one-hot encoding and *k*-mer-based approaches that directly reflect sequences. Moreover, these sequences are deficient in physicochemical characteristics, such as structural information and hydrophilicity; hence, they fail to accurately represent the inherent function of the given sequences. In this study, we utilized the physicochemical properties of amino acids to develop a rapid and efficient approach for extracting feature parameters that are suitable for downstream processes of machine learning, such as clustering. A fixed-length feature vector representation of a spike sequence with reduced dimensionality was obtained by converting amino acid residues into physicochemical parameters. Next, t-distributed stochastic neighbor embedding (t-SNE), a method for dimensionality reduction and visualization of high-dimensional data, was performed, followed by density-based spatial clustering of applications with noise (DBSCAN). The results show that by using the physicochemical properties of amino acids rather than conventional methods that directly represent sequences into numerical values, SARS-CoV-2 spike sequences can be clustered with sufficient accuracy and a shorter runtime. Interestingly, the clusters obtained by using amino acid properties include subclusters that are distinct from those produced utilizing the method for the direct representation of amino acid sequences. A more detailed analysis indicated that the contributing parameters of this novel cluster identified exclusively when utilizing the physicochemical properties of amino acids significantly differ from one another. This suggests that representing amino acid sequences by physicochemical properties might enable the identification of clusters with enhanced sensitivity compared to conventional methods.

**Author summary:** One of the major causes of the global threat of SARS-CoV-2 is the rapid emergence of its variants. While analyzing these variants is crucial for understanding the mechanism of outbreaks, the expansion of database size is becoming a barrier for effective analysis. In this study, we provide an approach that allows researchers without vast computational resources to comprehensively analyze the variants of SARS-CoV-2 spike by representing the sequences using the physicochemical properties of amino acids. The result of clusters derived using this method demonstrates not only an accuracy comparable to the conventional approaches of directly converting sequences into numerical values but also indicates the potential for more detailed clustering outcomes. The results suggest that our approach is valuable for the rapid identification of characteristic residues in new variants of SARS-CoV-2 and other viruses that may arise in the future.

## Introduction

The coronavirus disease 2019 (COVID-19) epidemic caused by severe acute respiratory syndrome coronavirus 2 (SARS-CoV-2) first began in Wuhan and went on to cause a massive global outbreak. One of the challenges in controlling the spread of this virus is the continuous appearance of variants [1–3]. Previous studies have shown that mutations in the spike protein, which binds to the human receptor hACE2, are associated with immune evasion and increased infectivity [4–6]. In addition, the combined effects of multiple mutations in the spike protein can largely affect infectivity and/or immune evasion. Variants with a single mutation at N501 were neutralized by antibodies, but a combined mutation of N501Y/K417N/E484K resulted in moderate resistance to antibodies [6]. Therefore, identifying and understanding these mutations will contribute significantly to reducing the risk of pandemics [7].

The infrastructure for depositing genomic and protein sequence data of these variants has been quickly established, allowing every researcher to access these data. As a result, over 3 million sequences of the variants of the SARS-CoV-2 spike protein have been registered on NCBI Virus. Even now, genome data sharing platforms such as NCBI Virus and GISAID are accumulating large amounts of amino acid sequences at a rate of several thousand or more per day [8, 9]. However, it is impractical to analyze all of them with the computational resources available to individual researchers, and it is challenging to identify and select important variants for *in vivo* or *in situ* analyses of mutations. Recent methods using machine learning, such as one-hot encoding [10] or *k*- mer-based methods [11], can be applied to the analysis of SARS-CoV-2 variants. However, the numerical representation of amino acid sequences using these methods raises concerns about the increase in dimensionality, which requires more computational resources and training data.

Furthermore, these methods have the disadvantage of not considering the physicochemical properties of amino acids when converting them into numerical variables, although recent studies have shown that changes in the physicochemical properties of amino acids, such as the hydrophobic shift in the N501Y mutation and the volumetric alteration in D614G, significantly influence infection function [3, 5]. Since the functions of proteins are determined by the physicochemical properties of amino acids and their arrangement, downstream analysis based on amino acid sequences, including machine learning approaches, should reflect the physicochemical properties of amino acids [12, 13].

In this paper, we propose a new approach for simple and rapid clustering of mutated spike proteins of SARS-CoV-2 variants with relatively small computational resources by using amino acid physicochemical properties as explanatory variables (Fig 1). Instead of converting the amino acid sequences into numerical values that represent the sequences themselves, we represented them by assigning volume and hydrophilicity parameters to the amino acid sequence of the spike protein. Utilizing these parameters, we successfully performed dimensional compression using t-SNE [14] and clustering with DBSCAN [15] and achieved a shorter runtime than that of methods employing one-hot encoding and the *k*-mer-based approach. Our method also facilitated the identification of novel subclusters and had the capability to identify important features, including the physicochemical information of amino acids.

**Fig 1.**
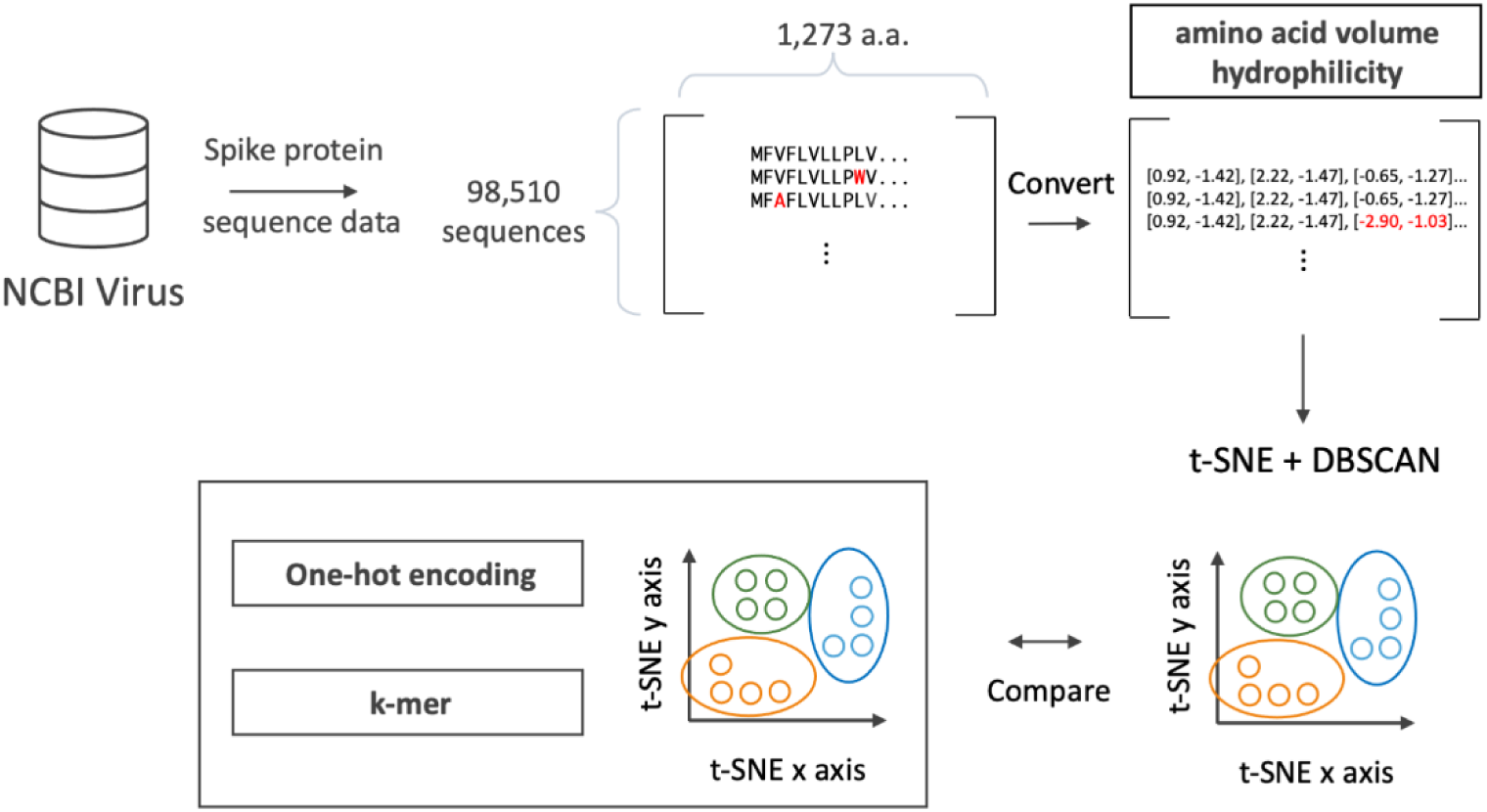
Schematic illustration of this study. First, the spike protein sequence of SARS- CoV-2 was downloaded from NCBI Virus. The obtained sequences were preprocessed to create a sequence × 1,273 amino acids (a.a.) matrix. These sequences in the matrix were then assigned volume and hydrophilicity values to represent amino acid parameters. The assigned data were dimensionally compressed with t-SNE and clustered with DBSCAN. The clustering result was compared to those of the one-hot encoding and *k*-mer-based method.

## Results

### Vectorization of the SARS-CoV-2 spike sequences

The dataset of spike protein sequences obtained from NCBI Virus was subjected to pairwise alignment while retaining the information of deleted residues (-) after removing sequences containing indeterminate residues (X, B, J, Z). The Wuhan type (WT) spike protein sequence was used as the reference sequence. After the pairwise alignments, sequences of spike proteins with an alignment length of 1,273 residues were collected. Details of this dataset are shown in S1 Table. Note that the alignment may contain residue deletions.

Next, each amino acid residue of the sequences in this dataset was converted by three methods: one-hot encoding and *k*-mer-based methods, both of which transform the sequence into categorical variables, and a method based on amino acid properties, in which we applied volume and hydrophilicity parameters to each amino acid residue (S2 Table). These two parameters are the relative volume and degree of hydrophilicity of each amino acid identified and used in previous studies [13, 16]. Thus, each sequence of the spike protein consisting of 1,273 residues was converted into data with sequential parameters of 26,733 dimensions for one-hot encoding, 9,261 dimensions for the *k*-mer-based method, and 2,546 dimensions for amino acid parameters. For positions with deleted amino acids, both values were set to zero in the amino acid parameters, while in the one-hot encoding and *k*-mer-based methods, it was represented as the 21^st^ character of a single amino acid.

### Dimensionally compression and cluster detection

The transformed sequence information was dimensionally compressed into a 2- dimensional space by applying t-SNE, a machine learning algorithm especially optimized for high-dimensional data, and plotted in Cartesian coordinates with coloring based on the variant classification by the World Health Organization (WHO). The plots obtained from the three methods were compared (Fig 2). In all three clusters, each variant with the same WHO label was clustered together in proximity, suggesting that two-dimensional parameters for a single amino acid are sufficient for clustering by t-SNE. Both the Delta and Omicron variants, which have numerous registered sequences, formed distinct groups with high similarity in the t-SNE coordinates, indicating the presence of multiple subpopulations within these variants. Note that variants categorized as “Omicron” according to the WHO label include a portion of the “other” variants in the cluster analysis. This suggests that part of the unclassified “other” category, which was introduced due to recent updates in the database, contains the “Omicron” variant. We summarized the number of data dimensions transformed by each method, as well as the corresponding t-SNE runtime (Table 1). The results revealed that the method utilizing amino acid properties significantly outperformed the other methods in terms of speed.

**Fig 2.**
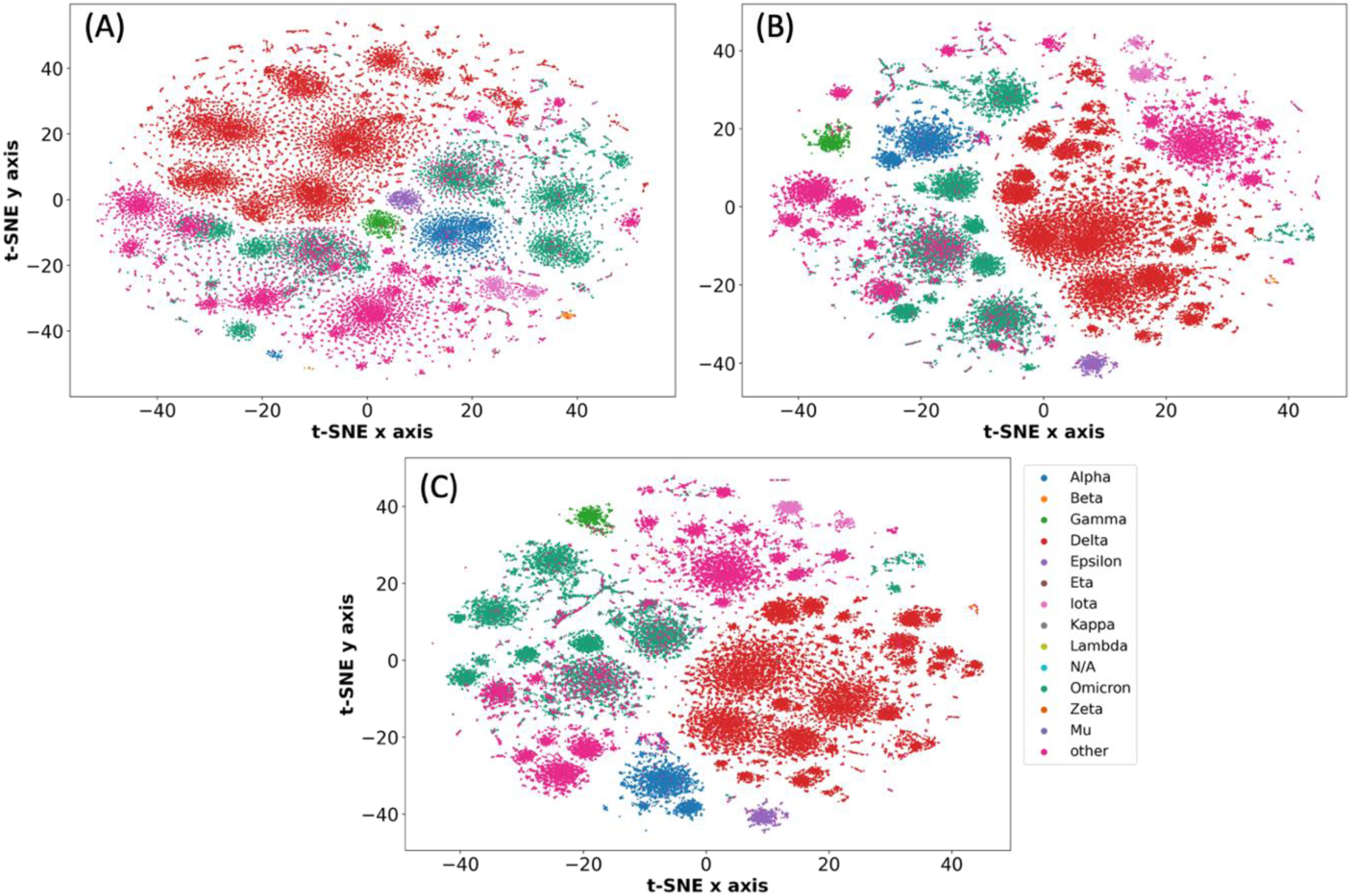
Dimensional compression and visualization of spike protein sequences by t- SNE. The sequences were converted into (A) the volume and hydrophilicity of each amino acid residue in the spike protein sequence, (B) the numerical values obtained by one-hot encoding and (C) the numerical values showing the frequency of each pair of three residues. Each value was dimensionally compressed by t-SNE. Each variant is shown in a different color.

**Table 1.**
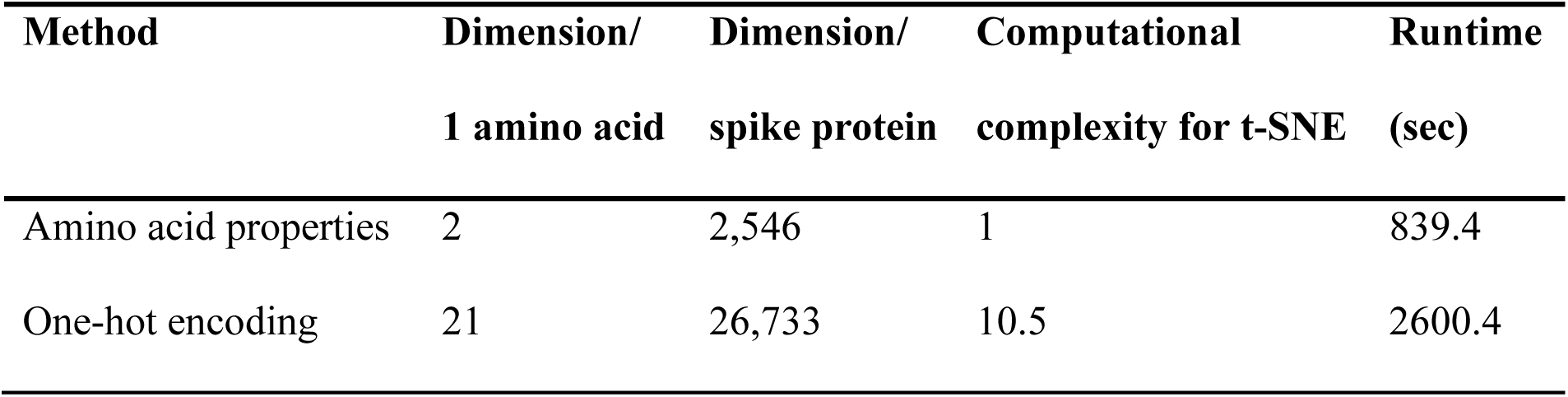
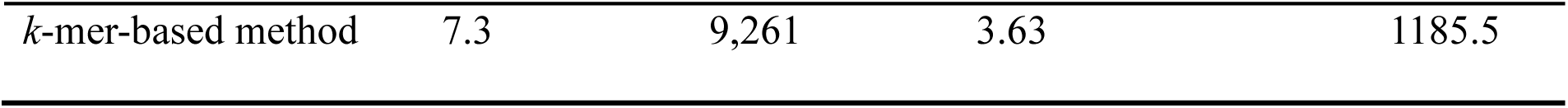
The number of dimensions and runtime for each method.

DBSCAN, a density-based clustering algorithm, was run to classify the data compressed by t-SNE. Clusters with sample sizes larger than the threshold (100 samples in this case) were extracted (Fig 3). The WHO-named variant that constitutes the largest population in each cluster and its proportion, along with its respective percentage, are shown in Fig 3. For example, a proportion of 100.0% means that the cluster contains only the specific WHO-named variant. We evaluated the clustering integrity by counting the number of clusters in which a single variant constituted more than 90% of the population, and 80 of 111 (72.1%) clusters determined by amino acid properties, 75 of 99 (75.8%) determined by one-hot encoding, and 78 of 95 (82.1%) determined by the *k*- mer-based method were considered to have clustering integrity. These results indicate that the three methods are comparable in the effectiveness of clustering.

**Fig 3.**
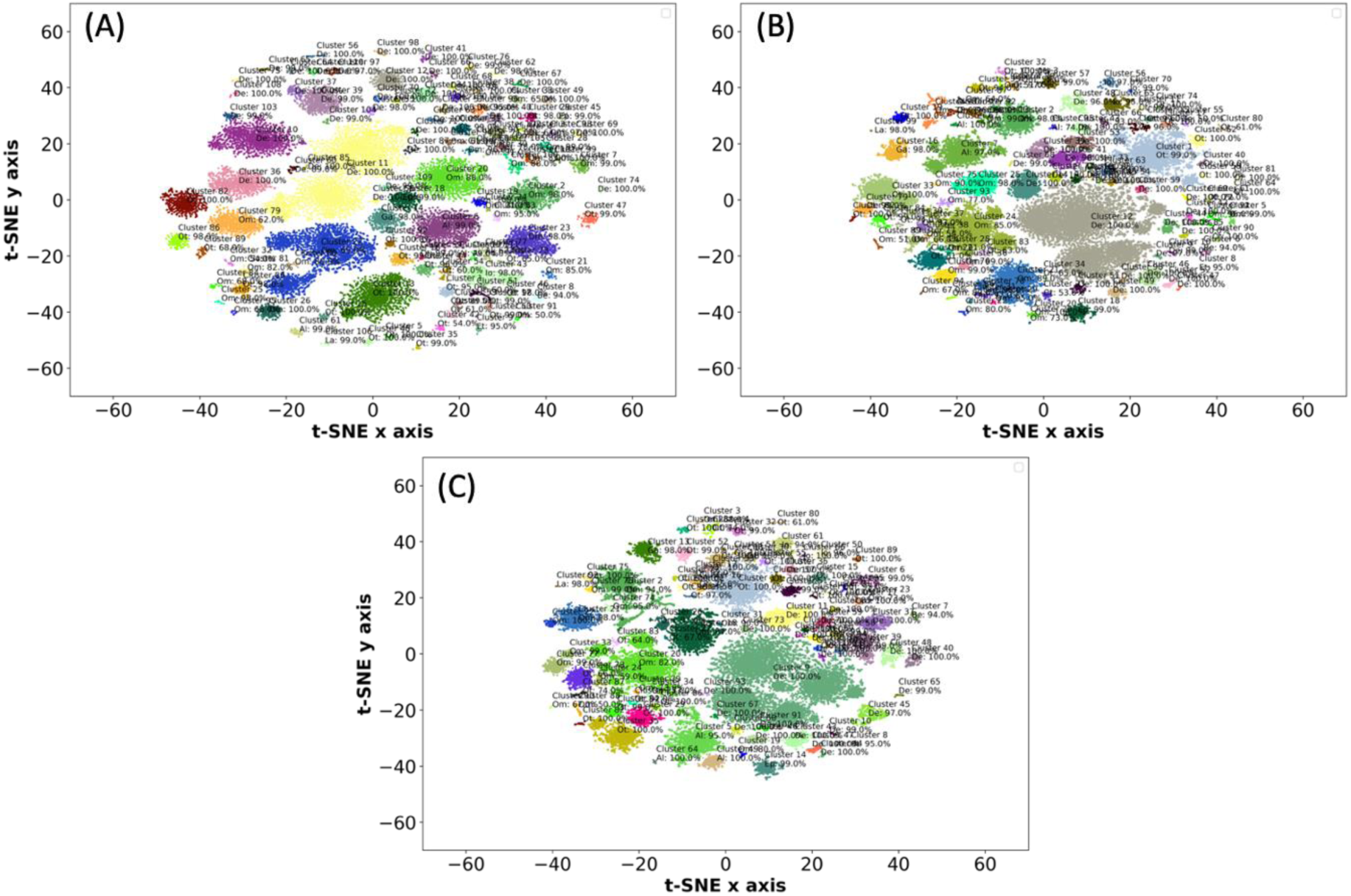
Clustering by DBSCAN. Each lineage was visualized by t-SNE, and the plot was divided into (A) 111 clusters in amino acid properties, (B) 99 clusters in one-hot encoding and (C) 95 clusters in the *k*-mer-based method. The labels indicate the cluster number, the most dominant variant in that cluster, and the percentage of the variant in the cluster. For example, the light-yellow cluster in the center of the coordinates, “Cluster 11 De: 100%,” means that it is the 11th cluster out of 111 clusters, the variant forming that cluster is Delta, and its percentage is 100%.

Additionally, to validate the utilization of amino acid properties as explanatory variables for evaluation of the spike of SARS-CoV-2, three classifiers (logistic regression, decision tree, and random forest) were applied to perform multiclass classification on each population of WHO-labeled variants (Table 2). The performance of each classifier was evaluated using five cross-validation runs. To compare the performance of each classifier by five cross-validations, we used the accuracy, precision, recall, and F1 score metrics. Accuracy represents the percentage of correctly classified cases in machine learning. Precision, also called the positive predictive value, describes the percentage of true positives among the positives determined by machine learning. Recall, also known as sensitivity, is the percentage of positives determined to be positive by machine learning out of all the positives. Since precision and recall have a trade-off relationship, the F1 score was used to evaluate their balance (see Methods). We found that all classifiers received high scores (Table 2), indicating that the WHO label was correctly predicted by using these amino acid properties obtained from the sequence of each variant. These results indicate the effectiveness of our method.

**Table 2.** Classification of clusters obtained by t-SNE.

|  | **Logistic Regression** | **Decision Tree** | **Random Forest** |
| --- | --- | --- | --- |
| Accuracy | $0.9335 \pm 0.0045$ | $0.9508 \pm 0.0013$ | $0.7801 \pm 0.0063$ |
| Precision | $0.9304 \pm 0.0351$ | $0.8871 \pm 0.0220$ | $0.6738 \pm 0.0179$ |
| Recall | $0.9317 \pm 0.0333$ | $0.9024 \pm 0.0250$ | $0.6766 \pm 0.0185$ |
| F1 score | $0.9369 \pm 0.0313$ | $0.9300 \pm 0.0369$ | $0.8840 \pm 0.0378$ |

Three classifiers, logistic regression, decision tree, and random forest, were used to classify each variant for the mappings reclassified by DBSCAN. The performance was evaluated by calculating the means and standard deviations of the accuracy, precision, recall and F1 score metrics for each 5-fold cross-validation trial. Data are expressed as the mean ± SD.

### Detailed comparison of clusterings by each method

To quantitatively compare the clusters obtained by each method, the similarity of the clustering results obtained through each method was evaluated by the adjusted Rand index (ARI) [17] and normalized mutual information (NMI) [18] (Fig 4). ARI is a quantitative indicator that evaluates the similarity between clusterings by each method by counting pairs assigned to identical or distinct clusters. An ARI score of 1.0 indicates completely identical clustering, while a score of 0 suggests random assignments. NMI is another indicator that quantifies the mutual information between two clusterings, normalized by the geometric mean of their individual entropies. A value of 1 indicates a perfect correlation between two clusterings, while a score of 0 denotes no correlation.

**Fig 4.**
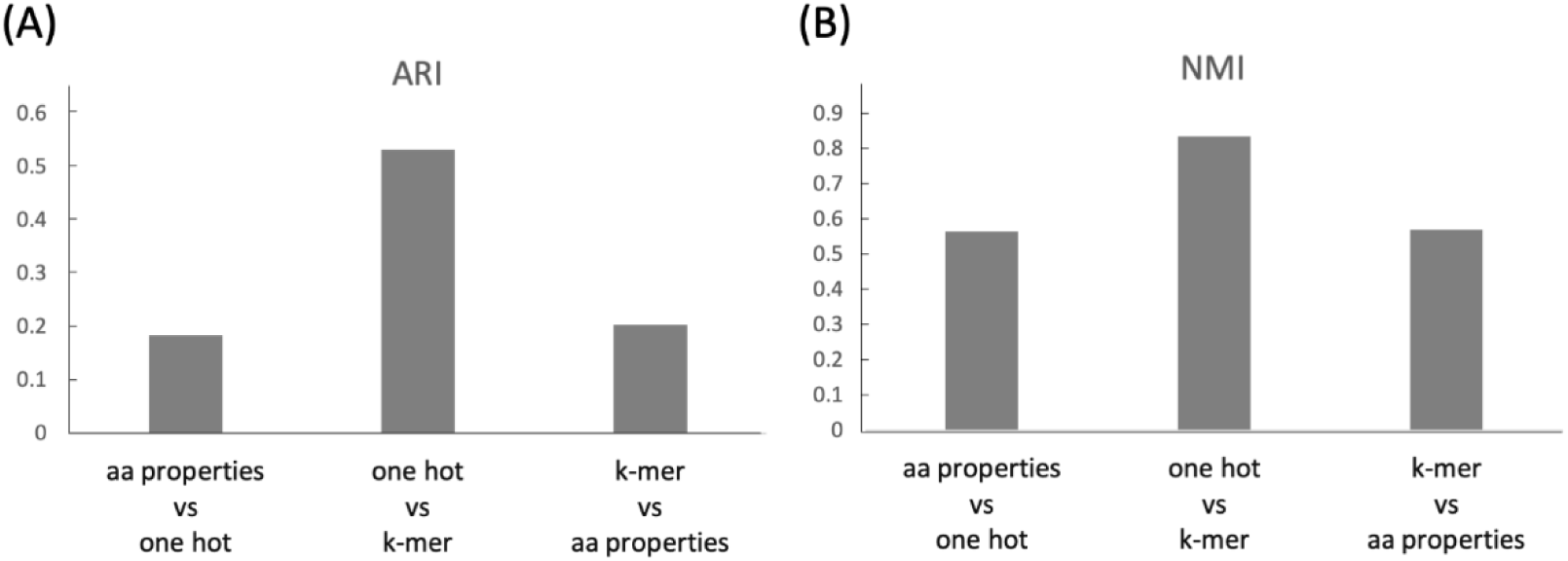
Comparing the similarity between each cluster. (A) The adjusted Rand index (ARI) for different clusterings by each method. The range of ARI values extends from - 1 to 1, where a value of 1 suggests perfect agreement between cluster assignments, a value of 0 suggests random clustering, and a value of -1 indicates a complete lack of correspondence between the two partitions. (B) Normalized mutual information (NMI) for different clusterings by each method. NMI values range from 0 to 1. A value of 1 indicates perfect information overlap between the two clusterings by each method, while a value of 0 indicates that the clusterings are entirely unrelated. aa properties: amino acid properties.

Notably, the similarity between the clustering results by one-hot encoding and the *k*- mer-based method demonstrated a high degree of similarity, while the similarity against clustering outcomes using amino acid properties was comparatively low. This lower similarity suggests that utilizing amino acid properties for clustering creates distinct clusters that reflect different features, capturing insights not accessible with sequence information alone.

To elucidate the specific differences in the clustering results obtained by these methods, we also compared the number and position of clusters for each WHO-labeled variant obtained by each method (S1-S3 Figs). As we expected, we found that clusters generated using amino acid properties differed significantly from those generated using other methods for certain labels. For example, when using amino acid properties, the Epsilon variants were divided into multiple distinct clusters (Fig 5). In contrast, with one-hot encoding and the *k*-mer-based method, only a single cluster of Epsilon was identified. Furthermore, these two Epsilon variant clusters detected with the amino acid properties were characterized using the random forest algorithm (Fig 6). Random forest is capable of identifying distinctive elements from each cluster. It revealed the location of the amino acid residues and the parameters that contribute to characterizing each cluster. The results revealed that W258 was the most contributing characteristic residue in one of the two Epsilon clusters. This differed significantly from another Epsilon cluster, despite both clusters belonging to the same Epsilon variant. In various cases, including the Epsilon case, there were significant differences in the characteristic residues within subclusters that were categorized as the same label. This result suggests that our set of methods provides a more detailed classification compared to one-hot encoding and *k*-mer-based methods.

**Fig 5.**
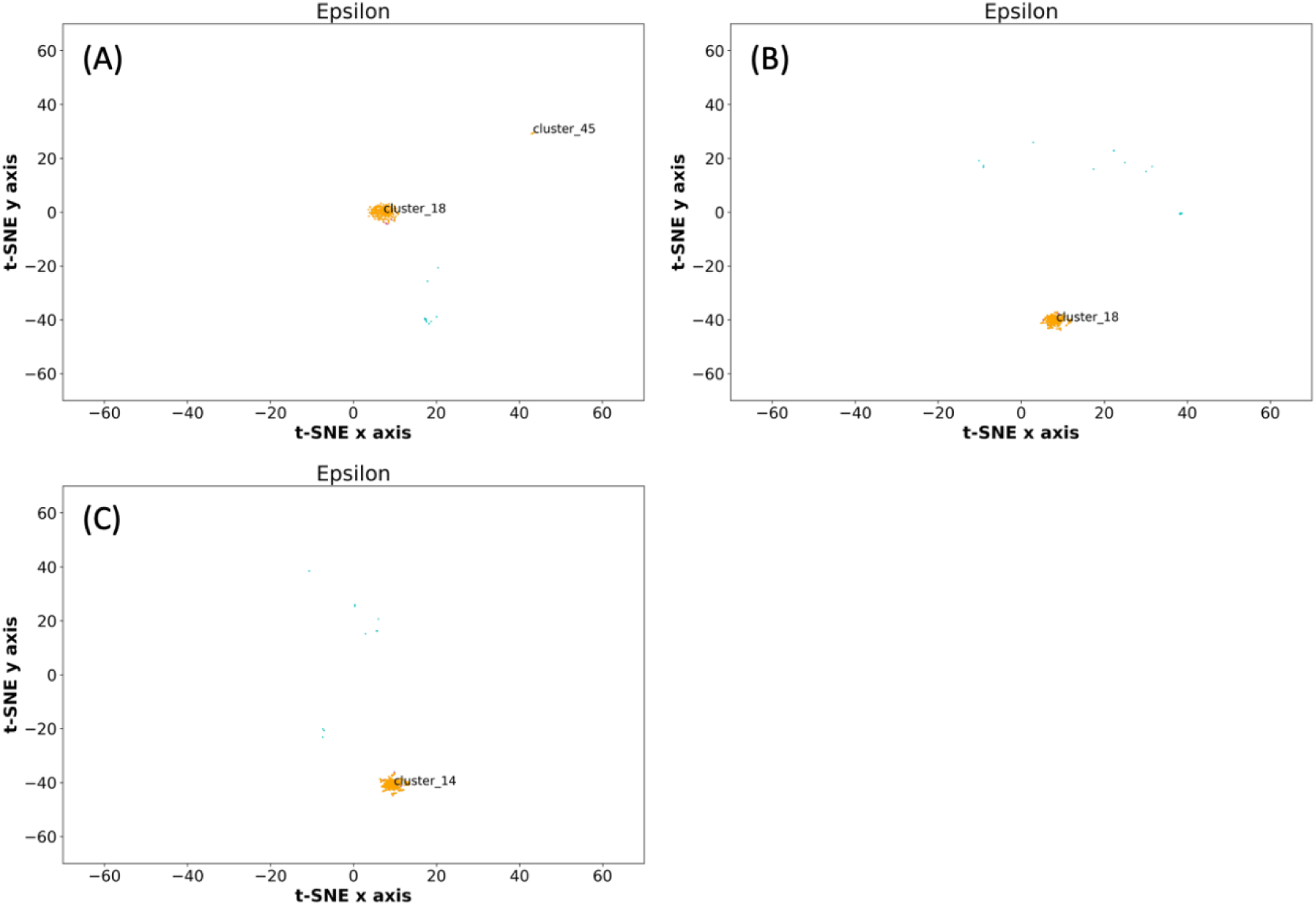
Comparing the Epsilon cluster obtained by each method. This figure is extracted from the result of DBSCAN shown in (A) Fig 3A obtained with amino acid properties, (B) Fig 3B obtained with one-hot encoding and (C) Fig 3C obtained with the *k*-mer-based method.

**Fig 6.**
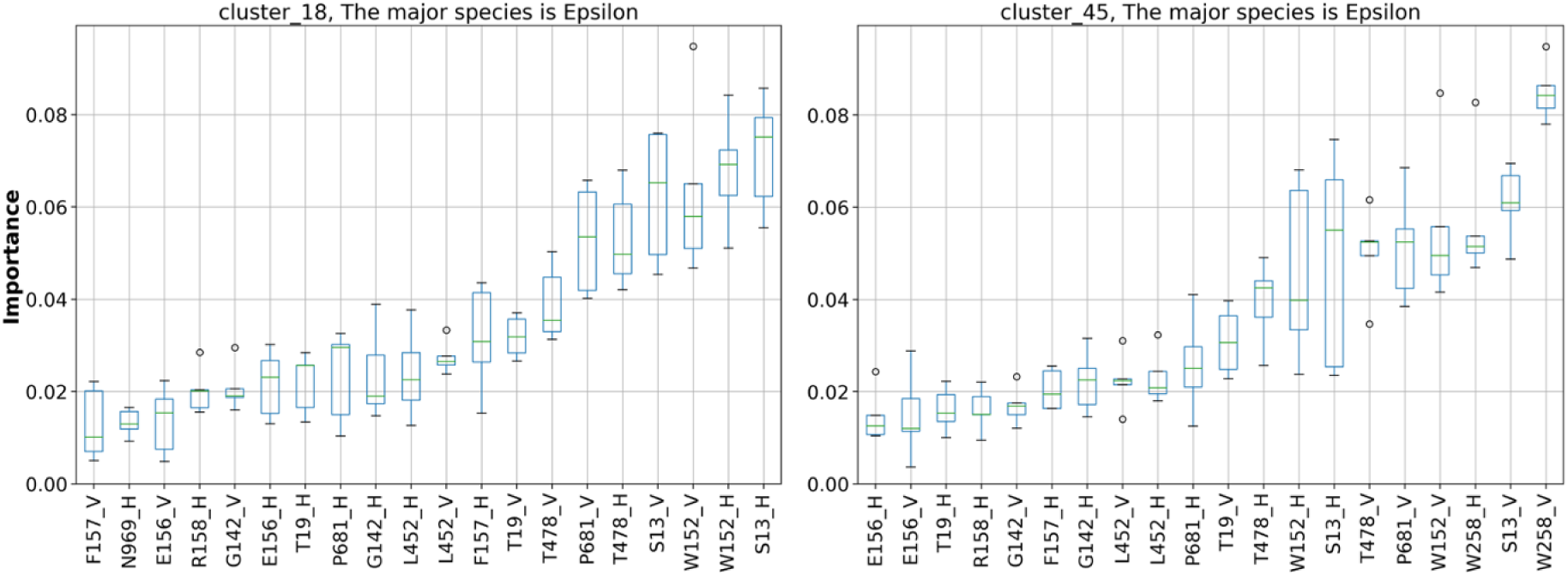
Characteristic residues in two Epsilon clusters detected with amino acid properties. Based on the clusters classified by DBSCAN, the top 20 characteristic residues and their physicochemical parameters were extracted by random forest. The hydrophilicity and volume of the amino acids are denoted by “H” and “V”, respectively. The horizontal axis indicates the amino acid residues of the spike protein and their physicochemical parameters. The vertical axis indicates the contribution of the physicochemical parameters of the corresponding amino acids to the random forest classification.

## Discussion

In this study, we introduced a rapid and efficient approach for clustering large sets of amino acid sequences based on their physicochemical properties. The clustering of spike protein variants represented by amino acid inherent physicochemical properties demonstrated sufficient accuracy compared to established methods such as one-hot encoding and *k*-mer-based methods, despite utilizing low-dimensional information. Furthermore, by utilizing physicochemical properties, we identified novel clusters and features that could not be classified by sequence-based methods, such as one-hot encoding and *k*-mer-based methods.

In the representation of amino acid residues in the method proposed in this paper, sequences are represented in a lower dimension (1,273 × 2 = 2,546) than in the conventional methods of one-hot encoding (21 × 1,273 = 26,733) and the *k*-mer-based method (21^3^ = 9,261), which reduces the dimension of the search space and the risk of the curse of dimensionality due to multidimensional analysis [19]. Remarkably, despite the low dimensionality, our clustering method accurately categorized WHO-labeled variants and correctly predicted each variant using only this low-dimensional parameter of amino acid properties. This suggests that this approach will be highly effective for managing the increasing volumes of data that are expected to increase as technology advances.

Previously, Kim et al. successfully reclassified the variants of SARS-CoV-2 spike using t-SNE and DBSCAN [20]. The methodology in that study transformed 3,822 gene sequences—corresponding to 1,274 amino acids multiplied by three—into a matrix with 3,721 dimensions. In contrast, our approach reduces this to a 2,548-dimensional matrix (1,274 amino acids × 2). Our methodology not only demonstrates superior computational efficiency and runtime but also provides insights into the physicochemical properties of amino acids, offering information that cannot be obtained through analysis by directly representing sequences. In fact, as shown in Fig 5, the cluster of the Epsilon variant was divided into two only when the physicochemical properties of amino acids were incorporated into the analysis. Furthermore, the two clusters generated based on amino acid properties exhibit residues with significant differences, suggesting that these clusters might represent biologically distinct variants (Fig 6).

Another feature of our method is its utilization of pairwise alignment. Because pairwise alignment allows sequences with different sequence lengths to be included in the dataset, this method can be applied even to variants with amino acid deletions. In addition, this method does not use multiple alignments, which are commonly used in phylogenetic tree analysis. The most time-consuming process in this method is dimensionality compression by t-SNE, and the computational complexity of t-SNE is O (LNlogN) [21], which is smaller than the complexity of multiple alignment (O(L^N^)) [22] in terms of the sequence length L and number of sequences N. Our method effectively reduces the computational cost and is easily adaptable to variations in data size as technological advancements emerge.

Whether *in vivo*, *in vitro*, or even *in silico*, detailed protein analysis requires substantial time and cost, limiting the number of protein samples that can be handled, and the selection of target proteins is an important issue [12, 23]. Because the method proposed in this paper assigns volume and hydrophilicity parameters that are considered important for protein function, the prediction results obtained by this method can serve as an indicator for selecting appropriate spike protein samples. In addition, the method proposed in this study can be used to identify characteristic amino acid properties at the residue level by evaluating the physicochemical similarity of the spike protein. Recognizing these key features of target residues enables the functional elucidation and modification of the spike proteins of the variants. Furthermore, the method proposed in this study can be customized and extended according to the objective by changing the physicochemical parameters to reflect the knowledge of researchers. Assigning other physicochemical parameters, such as charge, structure, and topology, may provide new insights into protein interactions [24].

## Conclusion

In this study, we developed an efficient and more in-depth unsupervised machine learning approach for SARS-CoV-2 spike analysis by utilizing amino acid physicochemical properties rather than employing raw sequence information. This method can be applied not only to SARS-CoV-2 spikes but also to other proteins if a sufficient sequence database is available. Beyond its utility for SARS-CoV-2 spikes, this method will provide an efficient way to address the analyses of a vast number of sequences in potential future pandemics.

## Materials and Methods

### Analysis environment and source codes

We performed this analysis using a MacBook Pro (16-inch, 2019, 2.4 GHz 8-Core Intel Core i9 processor and 64 GB 2667 MHz DDR4 memory). All analysis programs were coded in Python 3 and published on GitHub (https://github.com/horilab-dry/Covid_for_paper). Scikit-learn [25] was used to implement t-SNE, DBSCAN, random forest, ARI, NMI, logistic regression, and decision tree.

### Data collection and preprocessing

We obtained a sequence file of the spike protein from the NCBI Virus SARS-CoV-2 database (https://www.ncbi.nlm.nih.gov/labs/virus/vssi/#/virus?SeqType_s=Protein). We narrowed down the data to those that met the following criteria: “Proteins” was “surface glycoprotein”, “Sequence Length” was less than 1,273, and “Collection Date” was in the range of 10/1/2019-7/27/2023. When downloading the selected data, the fasta format was selected, and all records were specified to be downloaded. In the fasta definition line, “Accession”, “Pangolin”, “Host”, and “Collection Date” were selected. A one-to-one pairwise alignment was performed on all sequence data using the Wuhan type (WT). Sequence data containing irregular residues (X, B, J, Z) were deleted, and missing residues were assigned “-”. Only aligned sequences with a sequence length of 1,273 residues were extracted. To create the dataset for the downstream analysis, these sequences were converted into numerical values by using amino acid parameters, one-hot encoding and *k*-mer-based method. In amino acid parameters, the hydrophilicity and volume of the amino acids were assigned to each amino acid of the sequences. The volume and hydrophilicity parameters of the missing amino acids were set to 0. In the one-hot encoding and *k*-mer-based method, missing values “-” were converted to numerical values by handling them as single amino acids. This dataset contained 98,510 sequences. Details of the dataset are shown in Table 1.

### Data visualization with t-SNE

The t-SNE approach was implemented according to the original article [14]. t-SNE was applied to the characteristic parameters obtained by using amino acid parameters (volume and hydrophilicity), one-hot encoding and the *k*-mer-based method. In t-SNE, the parameter “perplexity” was used to determine the number of nearest neighbors among the data. In this study, all perplexity values were set to the default value of 30.

### Reclassification and performance evaluation with DBSCAN

DBSCAN was applied to the t-SNE result with two parameters, *ɛ* = 1.0 and *MinPts* = 30 [15]. Subsequently, the similarity of these clustering results obtained by t-SNE and DBSCAN was evaluated with the values of the ARI and NMI. Both the ARI and NMI were calculated between each method (amino acid properties vs. one-hot encoding, amino acid properties vs. *k*-mer-based method and one-hot encoding vs. *k*-mer-based method). The performance of the classifiers was evaluated by calculating the mean and standard deviation of the accuracy, precision, recall and F1 scores for each of the five cross-validation trials. Accuracy is an important measure that was used to assess the performance of the classification model. Accuracy was calculated as shown in the following equation:

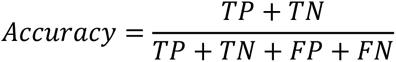

Precision is equal to the ratio of true positive (*TP*) samples to the sum of true positive (*TP*) and false positive (*FP*) samples. Precision is also a key metric to identify the number of correctly classified patients in an imbalanced class dataset. Precision was calculated by the following equation:

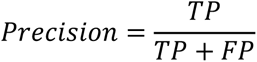

Recall is equal to the ratio of the true positive (*TP*) samples to the sum of true positive (*TP*) and false negative (*FN*) samples. Recall is an important metric for identifying the number of correctly classified variants in an imbalanced class dataset out of all the variants that could have been correctly predicted. Recall was calculated by the following equation:

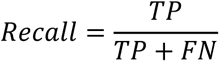

F1 score is equal to the harmonic mean of the recall and precision values. F1 score represents a balance between the precision and recall metrics and thereby provides a correct evaluation of the model’s performance in classifying variants. This is the most important measure that can be used to evaluate the model. F1 score was calculated by the following equation:

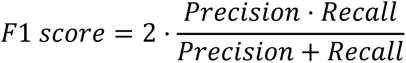

### Extracting and comparing the clusters of each WHO-labeled variant

The clusters of each WHO-labeled variant were extracted (S1-S3 Figs). Subsequently, the relative importance scores for each feature were calculated by a random forest algorithm [26]. The positions of amino acid residues with high contributions and the corresponding physicochemical parameters were identified. Random forest hyperparameters were run with n_estimators = 100 and max_leaf_nodes = 16.

## Acknowledgments

This work was supported by JSPS KAKENHI Grant Number JP21H05227 and AMED (20he0622029h0001).

## Author Contribution

**Conceptualization**: Takayuki Hashimoto, Katsutoshi Hori

**Funding acquisition**: Katsutoshi Hori

**Investigation**: Hiroya Oka, Kosaku Noba, Jun Sasahara, Takayuki Hashimoto

**Writing–original draft**: Hiroya Oka, Kosaku Noba, Jun Sasahara, Takayuki Hashimoto

**Writing–review & editing**: Shogo Yoshimoto, Hirohiko Niioka, Jun Miyake, Katsutoshi Hori

