## Supporting information for "Development of a fast feature extraction method for SARS-CoV-2 spike sequences using amino acid physicochemical properties"

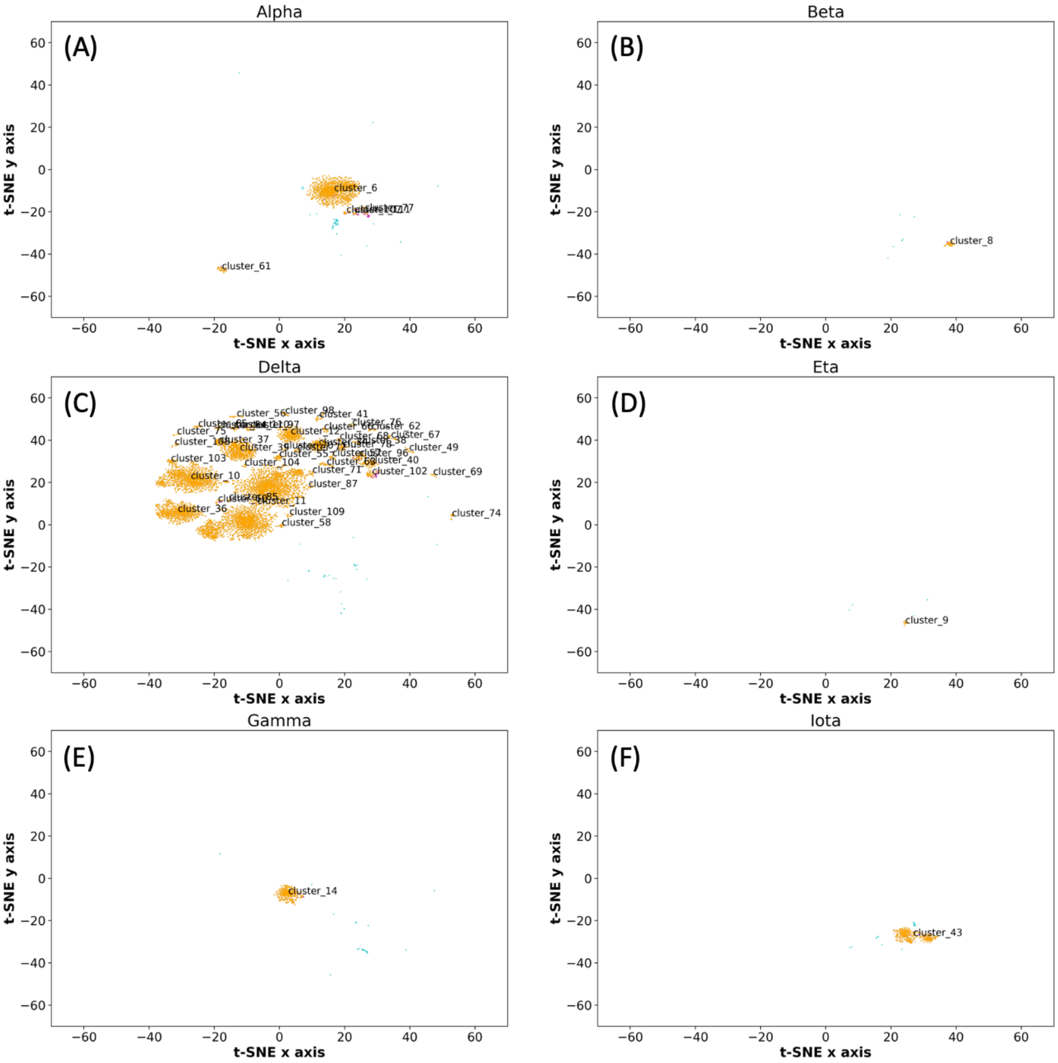
Alpha
(A)
cluster_6
cluster_61
Beta
(B)
cluster_8
Delta
(C)
cluster_56
cluster_41
cluster_75
cluster_37
cluster_103
cluster_104
cluster_10
cluster_87
cluster_11
cluster_36
cluster_109
cluster_58
cluster_49
cluster_67
cluster_96
cluster_40
cluster_71
cluster_102
cluster_69
cluster_74
Eta
(D)
cluster_9
Gamma
(E)
cluster_14
Iota
(F)
cluster_43
t-SNE x axis
t-SNE y axis

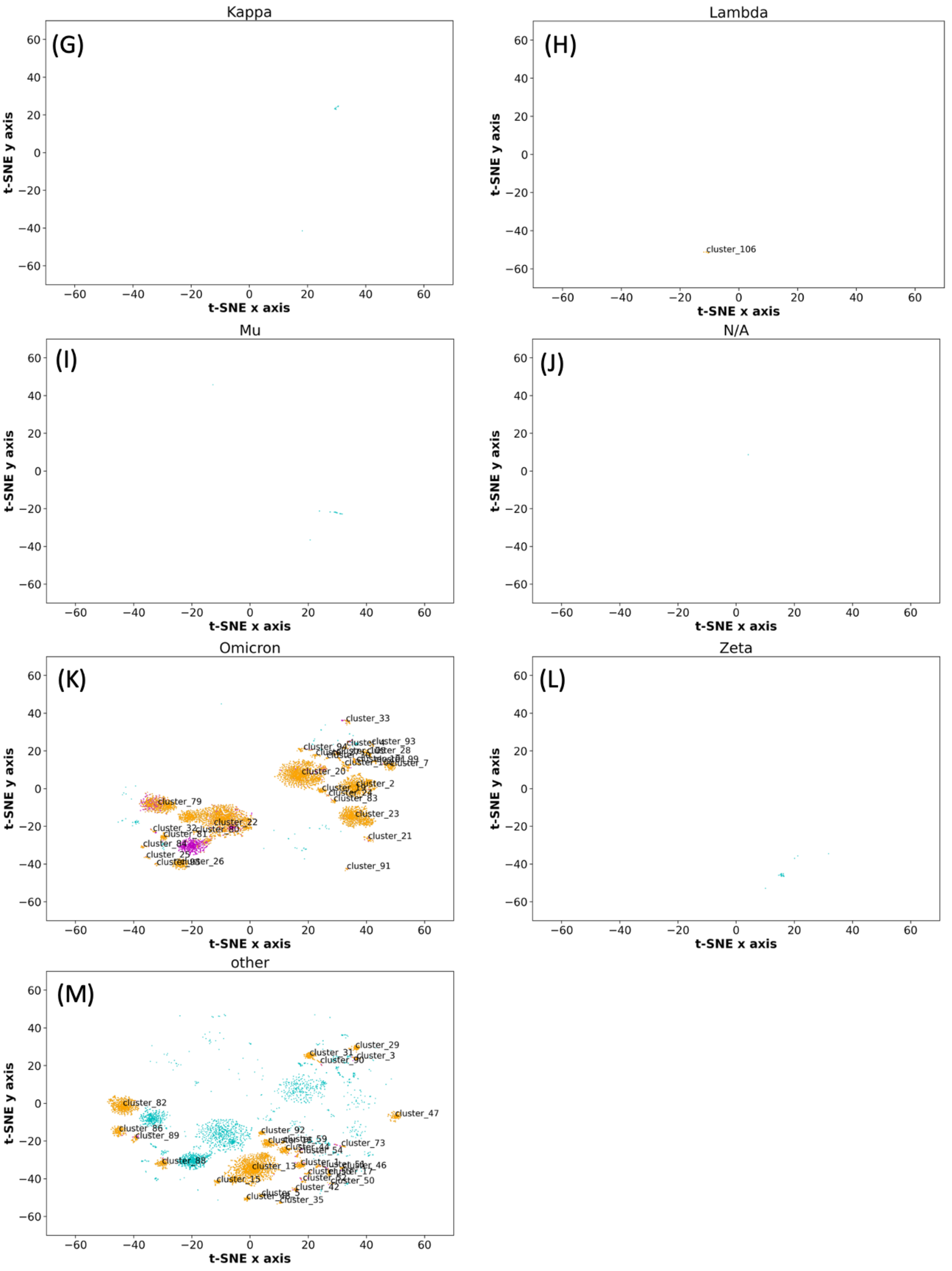


**S1 Fig. Dimensional compression by t-SNE and clustering by DBSCAN with amino acid properties, excluding the Epsilon variant.** The results shown in Fig 3A were extracted and displayed for each variant. Orange: The plot in which both the WHO label and detected cluster match the specified variant. Magenta: Only the DBSCAN-detected cluster matches the specified variant. Cyan: Only the WHO-specified label matches the specified variant. (A) Alpha, (B) Beta, (C) Delta, (D) Eta, (E) Gamma, (F) Iota, (F) Kappa, (H) Lambda, (I) Mu, (J) N/A, (K) Omicron, (L) Zeta, (M) Other.

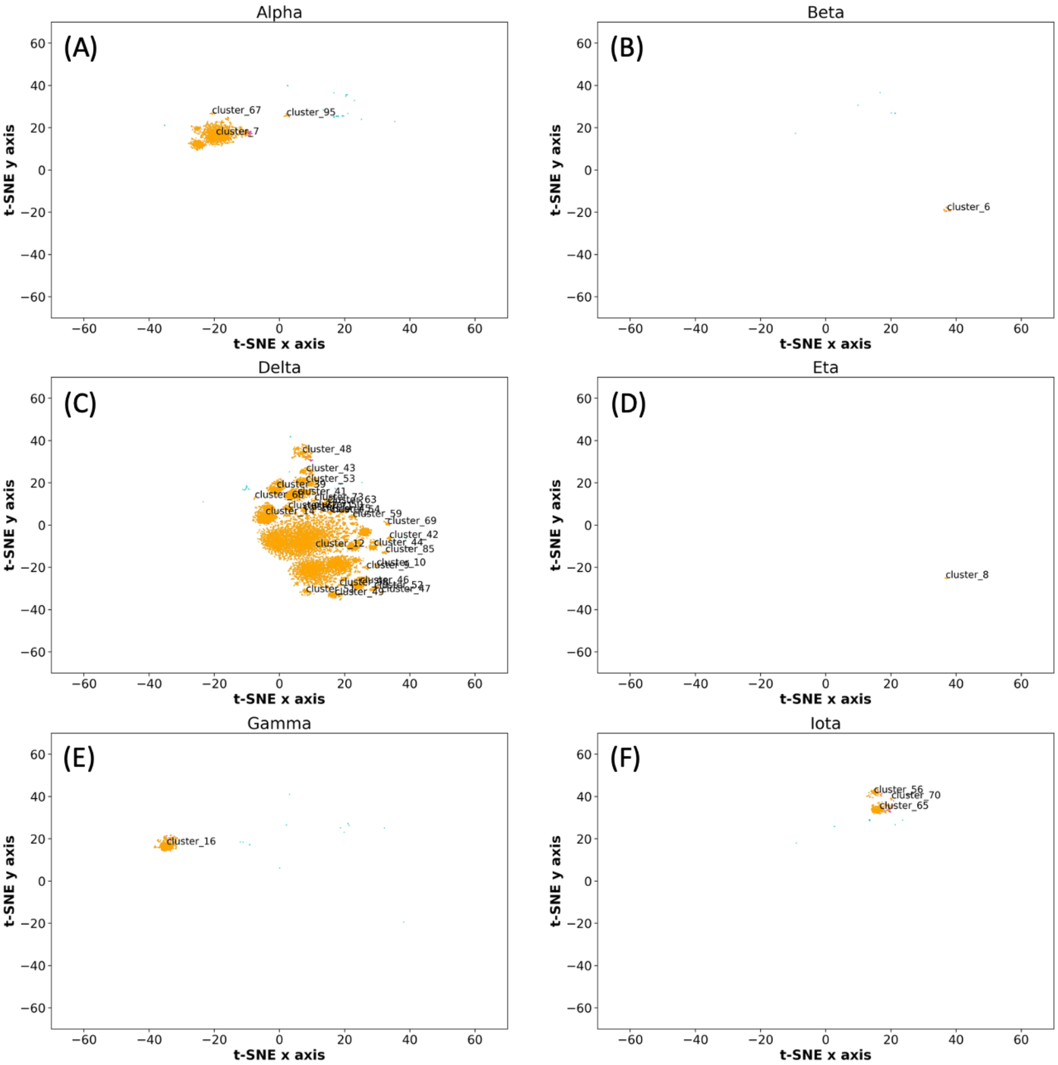
Alpha
(A)
cluster_67
cluster_95
cluster_7
Beta
(B)
cluster_6
Delta
(C)
cluster_48
cluster_43
cluster_53
cluster_69
cluster_42
cluster_12
cluster_85
cluster_10
cluster_47
Eta
(D)
cluster_8
Gamma
(E)
cluster_16
Iota
(F)
cluster_56
cluster_70
cluster_65
t-SNE x axis
t-SNE y axis

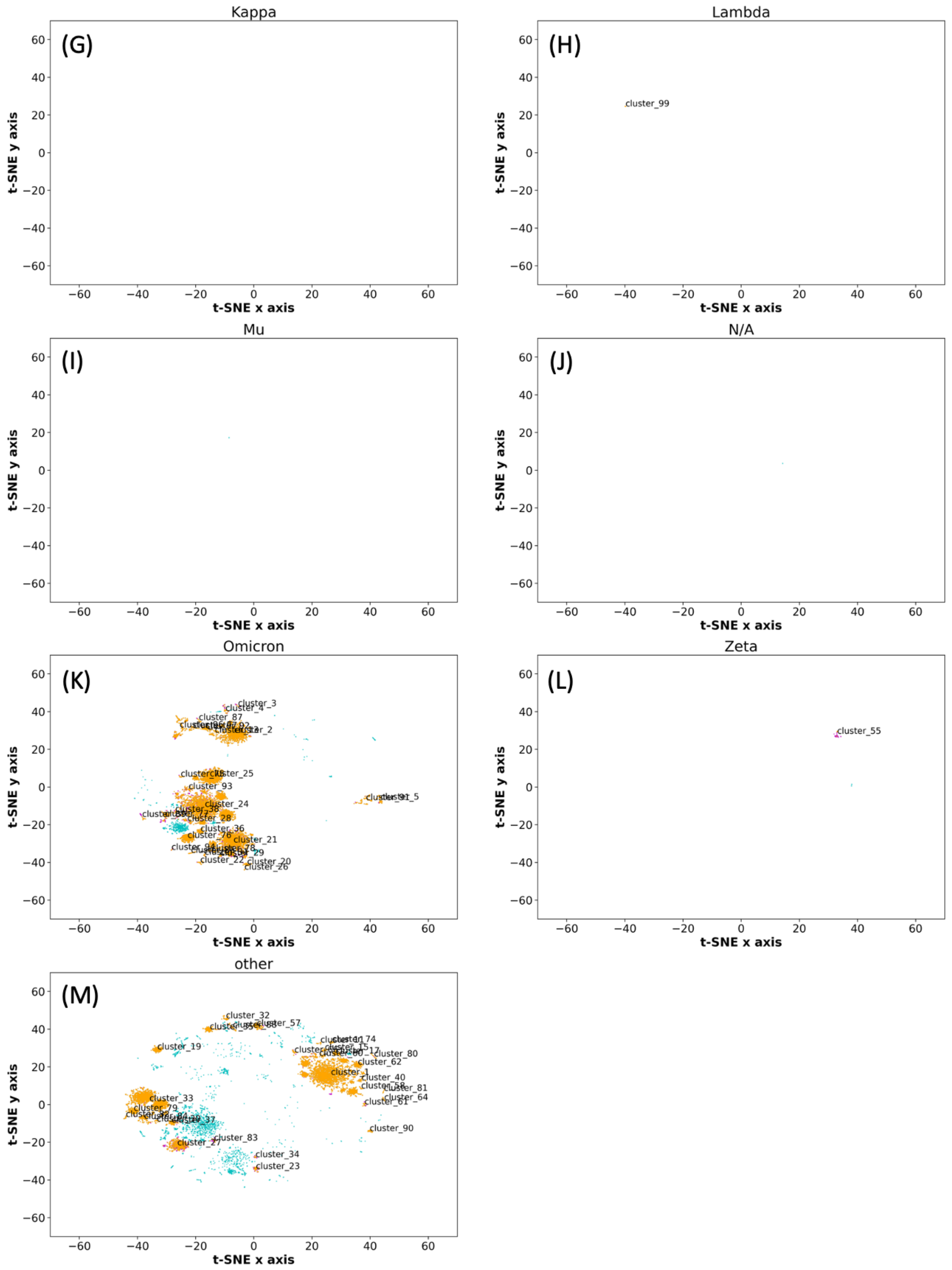


**S2 Fig. Dimensional compression by t-SNE and clustering by DBSCAN with one-hot encoding, excluding the Epsilon variant.** The results shown in Fig 3A were extracted and displayed for each variant. Orange: The plot in which both the WHO label and detected cluster match the specified variant. Magenta: Only the DBSCAN-detected cluster matches the specified variant. Cyan: Only the WHO-specified label matches the specified variant. (A) Alpha, (B) Beta, (C) Delta, (D) Eta, (E) Gamma, (F) Iota, (G) Kappa, (H) Lambda, (I) Mu, (J) N/A, (K) Omicron, (L) Zeta, (M) Other.

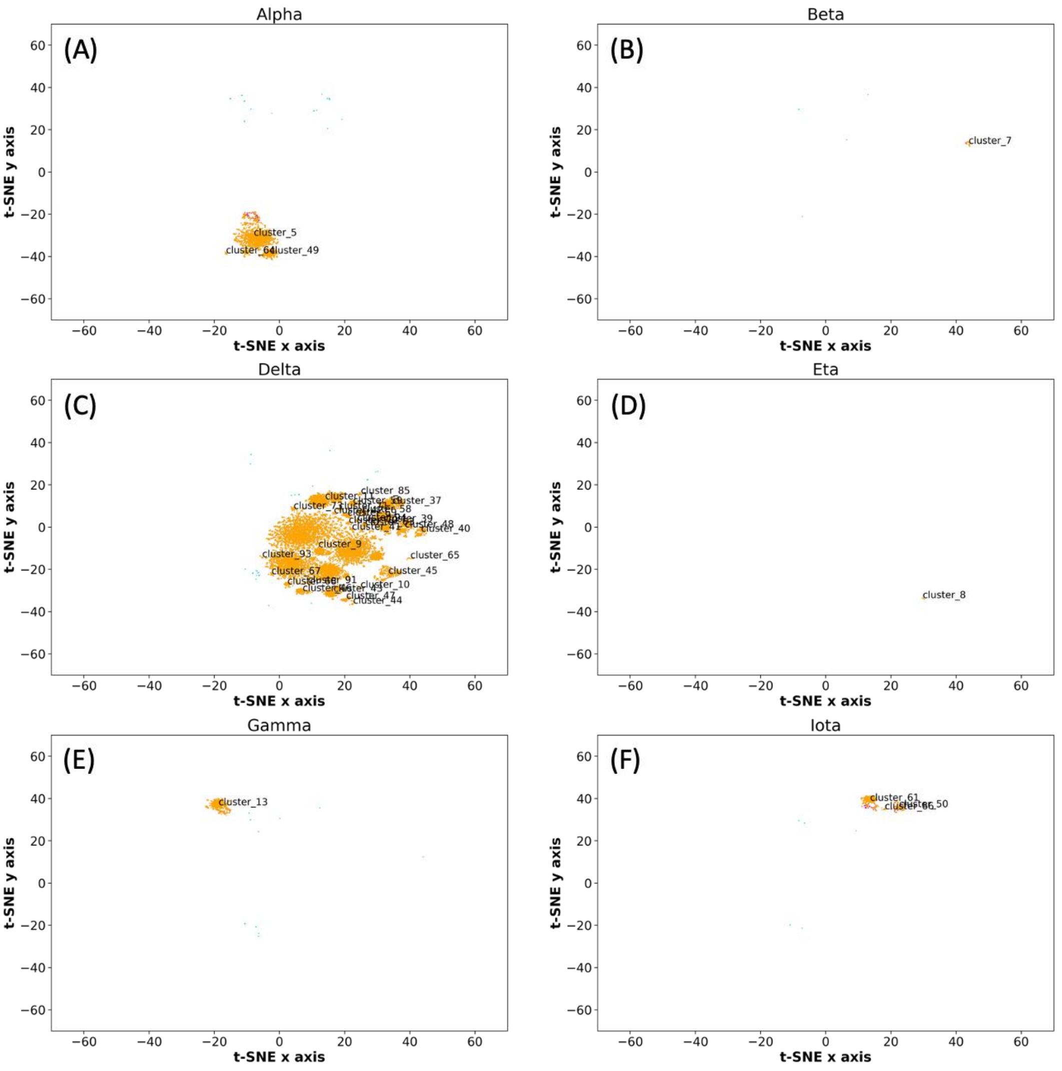
Alpha
(A)
cluster_5
cluster_64
cluster_49
Beta
(B)
cluster_7
Delta
(C)
cluster_85
cluster_37
cluster_39
cluster_48
cluster_40
cluster_9
cluster_93
cluster_65
cluster_67
cluster_45
cluster_10
cluster_47
cluster_44
Eta
(D)
cluster_8
Gamma
(E)
cluster_13
Iota
(F)
cluster_61
cluster_50
t-SNE x axis
t-SNE y axis

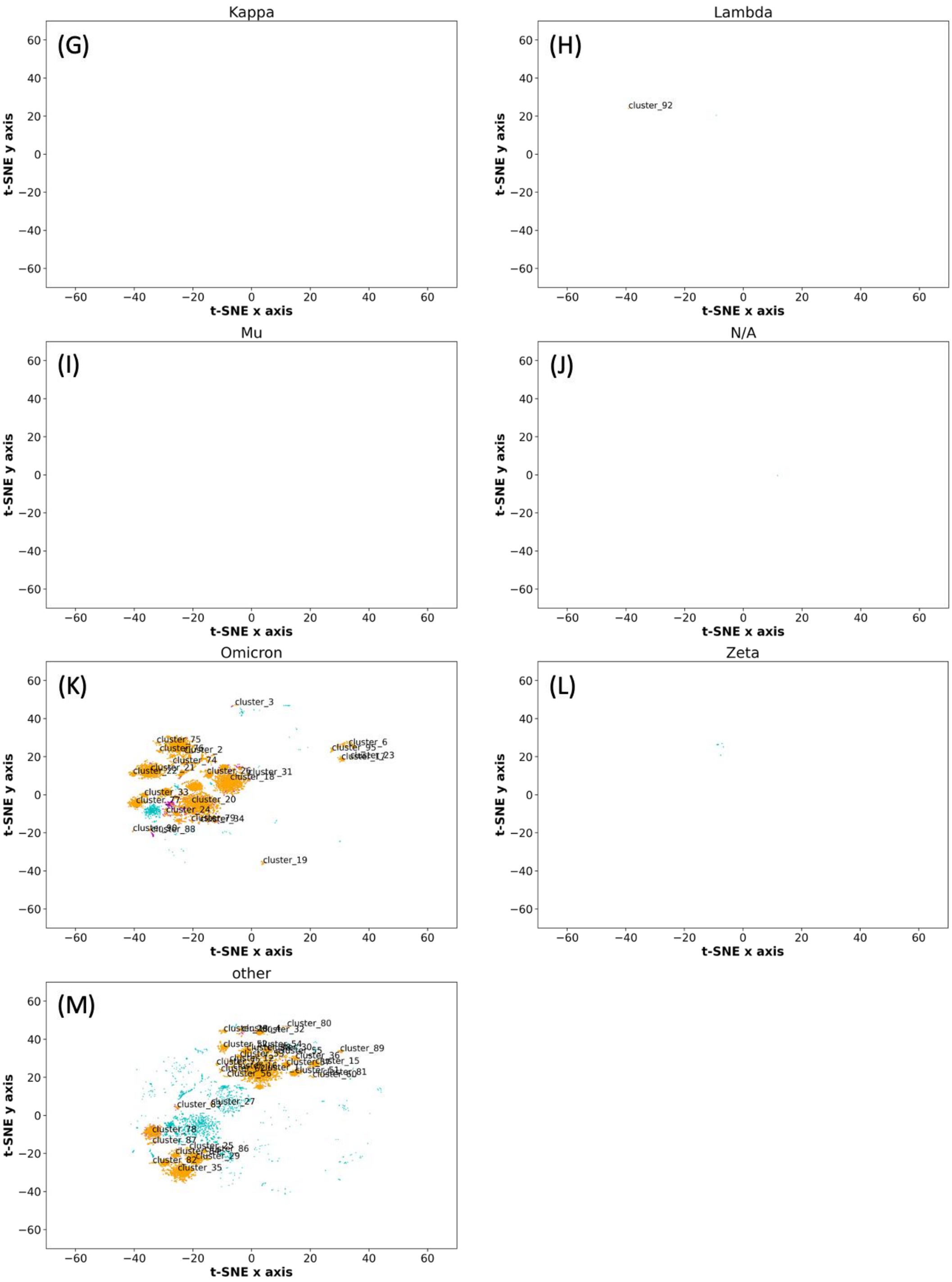


**S3 Fig. Dimensional compression by t-SNE and clustering by DBSCAN with the *k*-mer-based method, excluding the Epsilon variant.** The results shown in Fig 3A were extracted and displayed for each variant. Orange: The plot in which both the WHO label and detected cluster match the specified variant. Magenta: Only the DBSCAN-detected cluster matches the specified variant. Cyan: Only the WHO-specified label matches the specified variant. (A) Alpha, (B) Beta, (C) Delta, (D) Eta, (E) Gamma, (F) Iota, (G) Kappa, (H) Lambda, (I) Mu, (J) N/A, (K) Omicron, (L) Zeta, (M) Other.

**S1 Table. The correspondence between variants, their pangolin ID, and the numbers of sequences.** The number of sequences shown here is those that have been preprocessed and have a sequence length of 1,273. A total of 98,510 sequences were shown in the list and used for t-SNE, DBSCAN, and other processing.

| **Variant name** | **Pangolin** | **Number of sequences** |
|---|---|---|
| Alpha | B.1.1.7, Q* | 5,634 |
| Beta | B.1.351, B.1.351.* | 241 |
| Gamma | P.1.* | 1,542 |
| Delta | B.1.617.2, AY | 38,488 |
| Epsilon | B.1.427, B.1.429 | 1,316 |
| Eta | B.1.525 | 147 |
| Iota | B.1.526 | 1,606 |
| Kappa | B.1.617.1 | 40 |
| Lambda | C.37 | 132 |
| N/A | B.1.617.3 | 3 |
| Omicron | B.1.1.529*, XBB*, BA.*, XE | 24,877 |
| Zeta | P.2, P.3 | 121 |
| Mu | B.1.621, B.1.621.1 | 74 |
| Other | Other than those above | 24,289 |

**S2 Table. Volume and hydrophilicity parameters.** Each column represents one letter, the volume, and the hydrophilicity of the amino acid. Amino acid residues of deletions caused by local alignment were denoted by "-", and the respective parameter was set to 0.

| Amino acid (1 letter) | Volume | Hydrophilicity |
|---|---|---|
| A | -2.9 | -1.03 |
| C | -1.89 | 0.15 |
| D | -0.92 | 1.23 |
| E | 0.16 | 1.28 |
| F | 2.22 | -1.47 |
| G | -4.04 | 0.01 |
| H | 0.83 | 1.15 |
| I | 0.51 | -1.32 |
| K | 0.92 | 1.23 |
| L | 0.52 | -1.4 |
| M | 0.92 | -1.42 |
| N | -0.68 | 0.79 |
| P | -1.25 | -0.64 |
| Q | 0.36 | 1.09 |
| R | 2.41 | 1.31 |
| S | -2.36 | 0.38 |
| T | -1.19 | 0.28 |
| V | -0.65 | -1.27 |
| W | 4.28 | -0.18 |
| Y | 2.75 | -0.18 |
| - | 0 | 0 |
